# Dexamethasone-induced *PPARG* expression in osteogenic differentiation in vitro: impact on *SOX9* and *RUNX2* levels

**DOI:** 10.64898/2026.02.04.703935

**Authors:** Maria Rosa Iaquinta, Carmen Lanzillotti, Mauro Tognon, Fernanda Martini, Sonja Häckel, Martin J Stoddart, Elena Della Bella

**Author notes:** Corresponding author: Dr Elena Della Bella, Clavadelerstrasse 8, 7270 Davos Platz, Switzerland.

## Abstract

**Background:** The effects of dexamethasone during *in vitro* human osteogenesis present a complex picture. On one side, dexamethasone promotes the osteogenic differentiation of human bone marrow mesenchymal stromal cells (BMSCs) by downregulating *SOX9*. On the other side, it simultaneously promotes adipogenesis through the upregulation of *PPARG*. The regulation of *SOX9* and *PPARG* levels appears to be mediated by the transactivation function of the glucocorticoid receptor (GR), suggesting an indirect effect of dexamethasone on *SOX9* downregulation. This study aims to determine whether PPAR-γ affects the expression levels of *SOX9*, as suggested by several studies.

**Methods:** Human BMSCs were isolated from bone marrow and cultured in different osteogenic induction media containing 10 or 100 nM dexamethasone. Undifferentiated cells were used as control. Cells were treated either with a pharmacological PPAR-γ inhibitor (T0070907) or with a *PPARG*-targeting siRNA. Differentiation markers or PPAR-γ target genes were analysed by RT-qPCR. Mineral deposition was assessed by Alizarin Red staining. Two-way ANOVA followed by a Sidak multiple comparison test was used to compare the effects of treatments.

**Results:** Pharmacological inhibition of PPAR-γ had a mild effect on the expression of PPAR-γ target genes but hindered adipocyte formation. Neither *RUNX2* nor *SOX9* expression were affected by T0070907. siRNA treatment successfully downregulated *PPARG* expression, as well as that of PPAR-γ target genes *LPL, LPAR1*, and *ADIPOQ*. Contrary to expectations, *RUNX2* was significantly downregulated by the *PPARG*-siRNA treatment during osteogenic differentiation both in the absence and presence of dexamethasone, while *SOX9* levels were downregulated in undifferentiated cells. Overall, Alizarin Red staining analysis showed no change in mineralization levels when *PPARG* expression or activity was inhibited.

**Conclusions:** Understanding how dexamethasone regulates human BMSC differentiation is crucial to refine current *in vitro* models. These results suggest that PPAR-γ is not involved in *SOX9* or *RUNX2* repression during *in vitro* osteogenic differentiation of human cells.

## Background

Standard protocols for inducing *in vitro* osteogenic differentiation of human bone marrow mesenchymal stromal cells (BMSCs) largely depend on glucocorticoids and have remained mostly unchanged for decades. Dexamethasone, a synthetic glucocorticoid used clinically as an anti-inflammatory drug, is commonly combined with ascorbic acid to facilitate collagen production and a phosphate source to allow mineral formation (1).

A large body of literature suggests that dexamethasone enhances RUNX Family Transcription Factor 2 (*RUNX2*) expression and activity during *in vitro* osteogenic differentiation (2-4). However, in human BMSCs, *RUNX2* expression is largely unaffected by dexamethasone, with a high donor-to-donor variability. Its activity might be enhanced by the reduced expression of SRY-Box Transcription Factor 9 (*SOX9*), which has a dominant- negative effect on RUNX2 (5-7).

Dexamethasone binds with high affinity to the glucocorticoid receptor (GR). Upon ligand binding, GR induces a plethora of effects through genomic (both transactivation and transrepression of gene expression) and non-genomic events, the latter typically involving rapid mobilization of intracellular calcium (8). Notably, dexamethasone use for inducing osteogenic differentiation also leads to significant upregulation of Peroxisome Proliferator Activated Receptor Gamma (*PPARG*) expression and activity, resulting in the formation of adipocyte-like cells and adipokine production (6). This contrasts with the inverse relationship between bone marrow adipogenesis and osteogenesis, supporting the role of excess glucocorticoids in disrupting this balance (9-13).

Also, the mechanisms underlying dexamethasone and GR influence on *SOX9* expression need further investigation. Molecular dynamics simulations suggest a possible direct interaction between GR and SOX9 in the presence of dexamethasone, which might prevent SOX9 dimerization and its cis-acting effects (14). Although this mechanism might partially explain *SOX9* downregulation, we hypothesize that other factors are also involved. Previous data suggest that the effect of dexamethasone on *SOX9* downregulation is dependent on GR-mediated transactivation of gene expression, therefore indirect through an intermediate, upregulated target (6). Considering (a) the strong and dose-dependent regulation of *SOX9* and *PPARG* expression in opposite directions by dexamethasone, (b) the role of PPAR-γ in gene transrepression (15), and (c) reports of SOX9 and RUNX2 inhibition (13, 16, 17), it is plausible that PPARG is responsible for *SOX9* repression as a consequence of dexamethasone exposure.

To test this hypothesis, we inhibited either PPAR-γ activity or *PPARG* expression during dexamethasone-induced osteogenic differentiation. Pharmacological inhibition of PPAR-γ activity was achieved by treating human BMSCs with T0070907, a potent and selective PPAR-γ inhibitor (18). On the other hand, *PPARG* expression was transiently knocked down using a *PPARG*-targeting siRNA.

## Methods

### Cell isolation and expansion

The study was conducted in accordance with the Declaration of Helsinki. Human BMSCs were isolated as previously described (19, 20). Bone marrow was collected from patients undergoing spinal surgery with signed informed consent. Since the samples were anonymously collected, the Swiss Human Research Act did not apply and a full ethical approval was not required, as per the clarification of responsibility submitted to the cantonal authorities (Bern Req-2023-00198) (21). BMSCs were expanded in Minimum Essential Medium Eagle-Alpha Modification (α-MEM, Gibco, Thermo Fisher, Zürich, Switzerland), supplemented with 100 U/mL penicillin (Gibco), 100 _μ_g/mL streptomycin (Gibco), 5 ng/mL basic fibroblast growth factor (bFGF, Fitzgerald, Industries International, Acton, MA, USA), and 10% MSC-tested foetal bovine serum (FBS, Corning, USA). Cultures were maintained at 37°C, 5% CO_2_, 90% rH, and the medium was refreshed every second day. Cells were used at passage 3 for all the experiments.

### Inhibition of PPARG activity during osteogenic differentiation

BMSCs from 4 donors were used for this experiment (2M, 2F, median age 47 years, range 31-74). Osteogenic differentiation was induced as previously described (6, 20) in the presence of 50 _μ_g/ml L-ascorbic acid 2-phosphate sesquimagnesium salt hydrate and 5 mM β-glycerol phosphate (Sigma-Aldrich). Medium was further supplemented with either 10 or 100 nM dexamethasone-cyclodextrin complex (dex, Sigma-Aldrich). Cells maintained in growth medium (DMEM 1 g/l glucose, 10% FBS, 100 U/mL penicillin, 100 _μ_g/mL streptomycin) or without dexamethasone were used as controls.

PPAR-γ activity was inhibited by treating BMSCs with 10, 100, or 1000 nM T0070907 (Sigma-Aldrich), or using dimethyl sulfoxide (DMSO) as a vehicle control, in combination with the differentiation media.

Medium was refreshed three times/week. Samples were collected at day 7 for gene expression analysis (n=4 donors) and at day 21 for evaluation of mineral deposition (n=3 donors). T0070907 treatment was maintained constant for the whole duration of the experiment.

### Inhibition of PPARG expression in osteogenic differentiation

BMSCs from 4 donors were used for this experiment (3M, 1F, median age 68 years, range 50-85). Osteogenic differentiation was induced as described in the previous paragraph. A transient knock-down of *PPARG* expression was achieved by transfecting cells with 10 pmol/well of a *PPARG*-targeting siRNA (Silencer™ Select Pre-Designed siRNA, ID s10886, Ambion, Thermo Fisher) using the Lipofectamine™ RNAiMAX Transfection Reagent (Thermo Fisher). Cells that were not transfected, or transfected with the Silencer Select Negative Control No. 2 siRNA (siNeg), were used as controls. Transfection was performed concurrently with the first medium change, 24 hours after cell seeding, and was not repeated thereafter. Medium was refreshed three times/week. Samples were collected at day 7 for gene expression analysis and at day 21 for evaluation of mineral deposition (22).

### Alizarin Red staining and quantification

Mineral deposition was evaluated by Alizarin Red staining as previously described (6). Briefly, samples were fixed in 4% neutral buffered formalin and stained with an Alizarin Red S solution (40 mM, pH 4.2, Sigma-Aldrich). Following an extensive washing procedure, images were captured for qualitative evaluations. A quantitative analysis was performed with elution of the Alizarin Red from the stained samples, using the cetylpyridinium chloride method, as previously described (6). Eluate absorbance was measured at 540 nm using an Infinite® 200 PRO microplate reader (Tecan, Männedorf, Switzerland).

### RNA isolation and RT-qPCR

A day 7, cells were lysed in TRIreagent and used for RNA isolation using a phenol-chloroform extraction as previously described (6). Quantity and purity of RNA was evaluated using a Nanodrop One UV-Vis spectrophotometer (Thermo Fisher). TaqMan Reverse Transcription reagents (Applied Biosystems, Foster City, CA, USA) were used to synthesize cDNA from 500 ng of total RNA, as per manufacturer’s protocol. Gene expression was evaluated using a TaqMan Gene Expression Master Mix (Applied Biosystems) in a QuantStudio 6 Pro Real-Time PCR system (Applied Biosystems). Assay details are reported in **Table 1**. TaqMan gene expression assays (ThermoFisher) or custom-designed primers and probes (Microsynth AG, Balgach, Switzerland) were used. Relative quantification (RQ) of gene expression was calculated as 2^-ΔΔCt^ (endogenous control: *RPLP0*; calibrator: cells grown in growth medium, DMSO control or siNeg control).

**Table 1.**
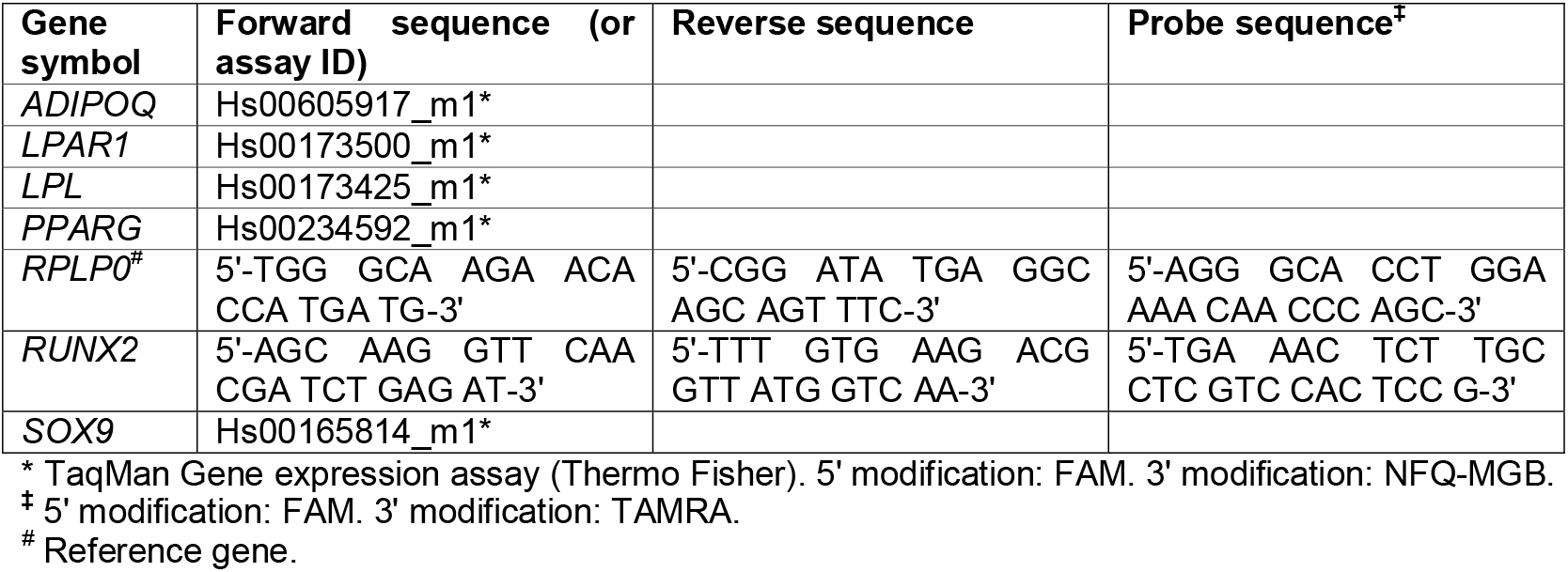
qPCR assay details.

### Statistical analysis

Statistical analysis was performed using GraphPad Prism v.10 (GraphPad Software, San Diego, CA, USA). A RM two-way ANOVA with Sidak‘s multiple comparison test was used to compare the effect of either the siRNA or T0070907 compared to the relative controls (siNeg and DMSO, respectively). Results were considered significant when *p* < 0.05.

## Results

First, different concentrations of T0070907 were screened in a preliminary experiment. The PPAR-γ inhibitor was tested at 10 nM, 100 nM, and 1 _μ_M. The highest concentration (1 _μ_M) had a detrimental effect on cell viability, as observed by morphological changes (Supplementary file, **Fig. S1**). Preliminary data suggested that using 100 nM T0070907 led to a more consistent PPARG inhibition (data not shown). Therefore, subsequent experiments were conducted using this concentration.

At 100 nM, treatment of human BMSCs with T0070907 mildly inhibited PPAR_-γ_ activity, as indicated by the non-significant downregulation of its targets *ADIPOQ* and *LPL* (**Fig. 1A**). *PPARG* expression was not significantly altered by T0070907 treatment but was upregulated by dexamethasone, as expected, along with *PPARG* target genes (**Fig. 1A**). By day 21, treatment with 100 nM T0070907 resulted in the absence of adipocyte-like cells, which are typically observed in monolayers cultured with 100 nM dexamethasone and, to a lesser extent, with 10 nM dexamethasone (**Fig. 1B**).

**Figure 1.**
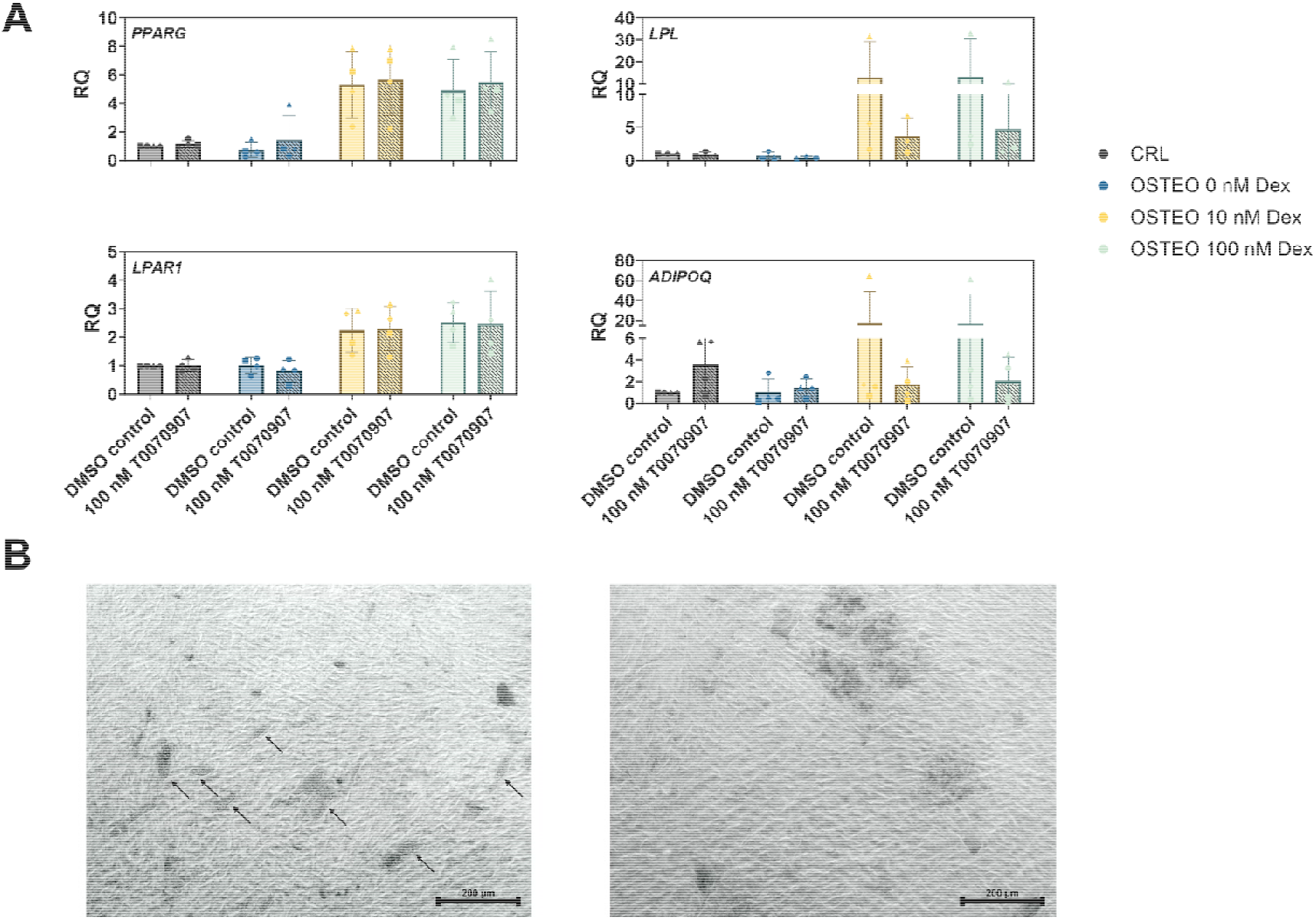
**A)** Relative quantification (RQ) of *PPARG* and PPAR-γ target genes expression at day 7. Solid filled bars: DMSO control. Striped bars: 100 nM T0070907. **B)** Brightfield, representative images of day 21 cultures from one donor. Left: cells cultured in the presence of 100 nM dexamethasone, DMSO control. Arrows indicate adipocyte-like cells. Right: cells cultured in the presence of 100 nM dexamethasone, treated with 100 nM T0070907. Scale bars: 200 μm.

At day 7, neither *RUNX2* and *SOX9* expression, nor their ratio, were affected by treatment with T0070907 (**Fig. 2A**). Alizarin Red staining did not show significant differences in mineral deposition after pharmacological inhibition of PPAR-γ activity (**Fig. 2B-C and S2**). The levels of osteogenic markers were mostly affected by dexamethasone.

**Figure 2.**
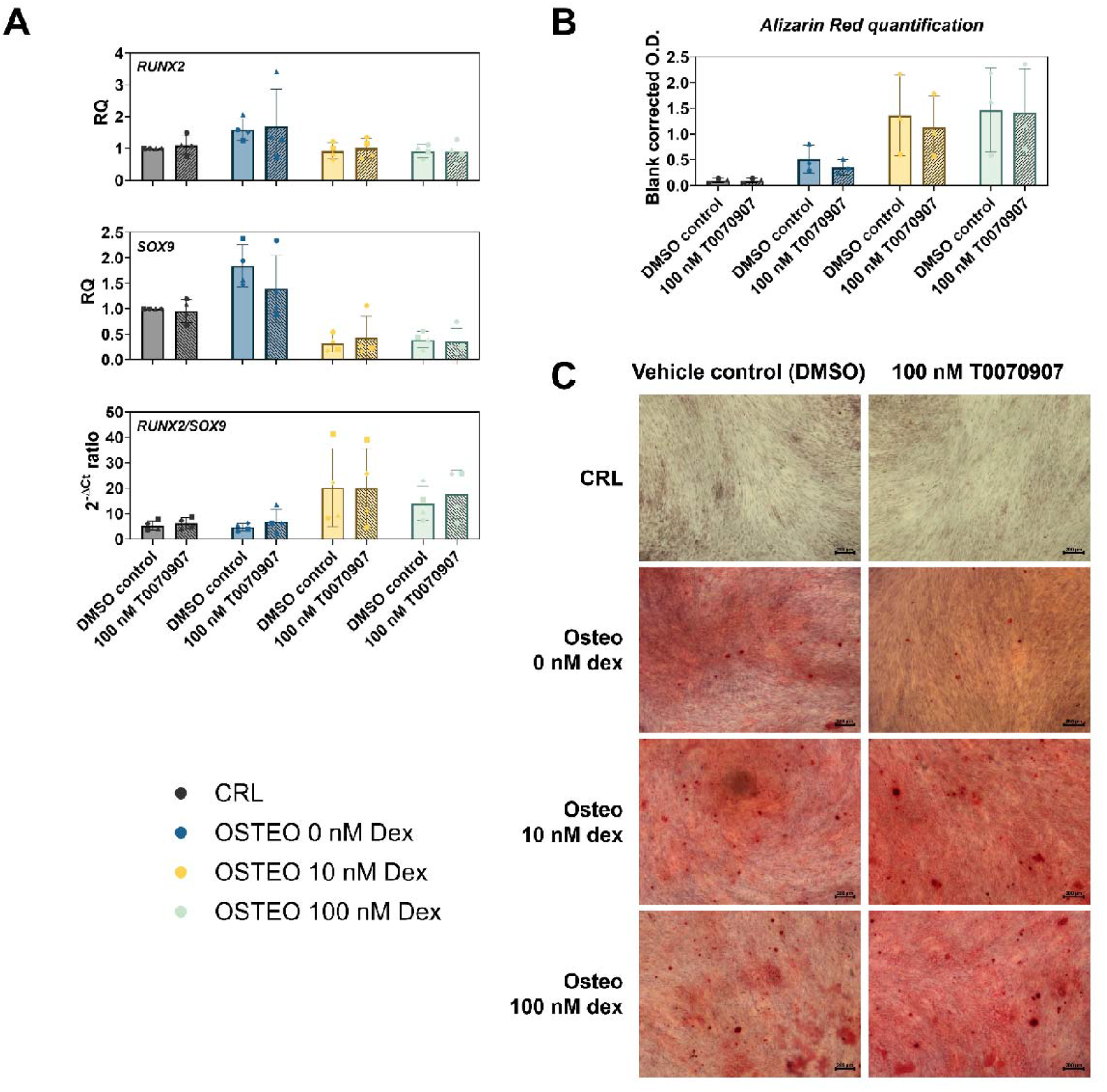
**A)** Relative quantification (RQ) of *RUNX2* and *SOX9* expression, and *RUNX2*/*SOX9* ratio levels, at day 7. Each symbol represents one donor (n=4). **B)** Alizarin Red staining quantification from n=3 donors and **C)** representative pictures from one donor at day 21 (scale bar = 200 μm). Solid filled bars: vehicle (DMSO) control. Striped bars: 100 nM T0070907.

The mild inhibition of PPAR-γ activity observed with T0070907 in our experimental conditions prompted us to transiently reduce *PPARG* expression through the application of a *PPARG*-targeting siRNA.

*PPARG* knock-down persisted at least for 7 days after initial siRNA transfection (**Figure 3**). At this time point, average *PPARG* downregulation after siRNA treatment was at least 10- fold compared to the siRNA negative control. Analysis of PPAR-γ transcriptional target expression revealed significant downregulation of *LPL, LPAR1* and *ADIPOQ* following *PPARG*-siRNA treatment in dexamethasone-containing groups (**Figure 3**).

**Figure 3.**
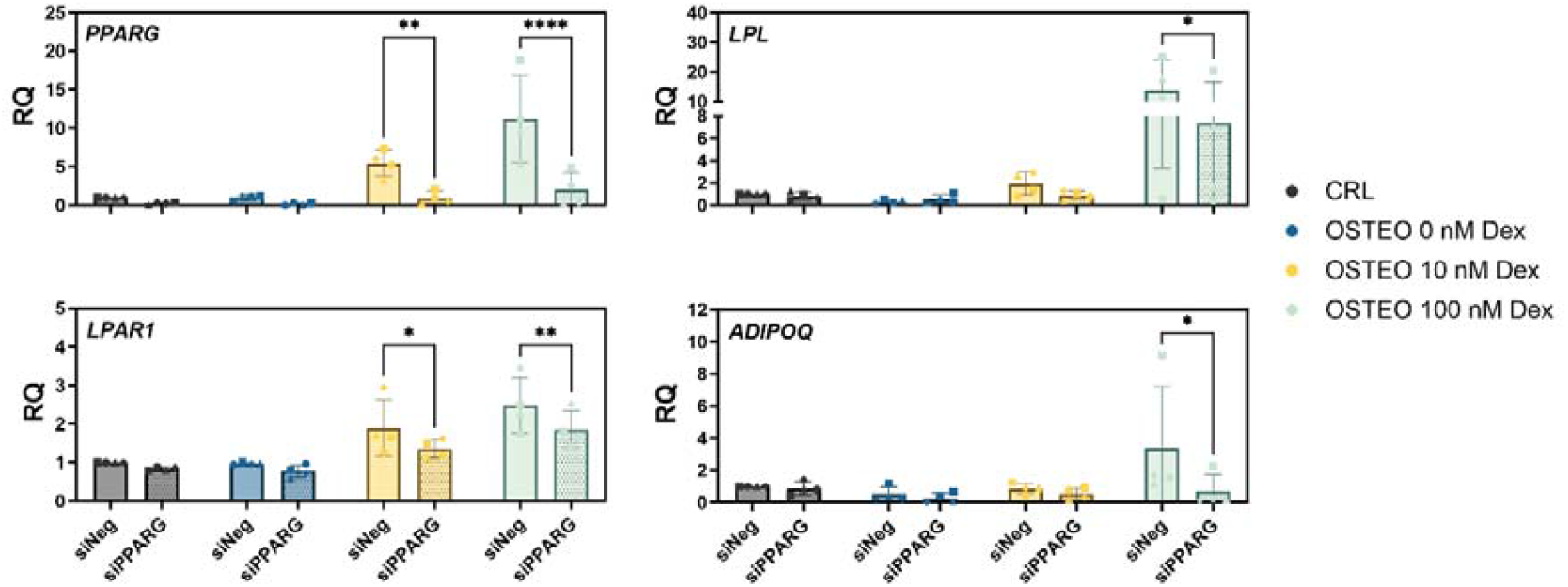
Relative quantification (RQ) of *PPARG* and *PPARG* target gene expression at day 7. * p < 0.05; **; p<0.01; **** p<0.0001. Solid filled bars: siRNA negative control. Dotted bars: treatment with *PPARG* siRNA.

Dampening of *PPARG* expression and activity was associated with either no effects or to a downregulation in both *RUNX2* and *SOX9* expression at day 7 (**Figure 4A**). These changes did not translate into significantly different *RUNX2*/*SOX9* ratios between the siNeg and si*PPARG* groups, which is consistent with the absence of effects of si*PPARG* treatment on mineral deposition at day 21 (**Figure 4B-C and S3**).

**Figure 4.**
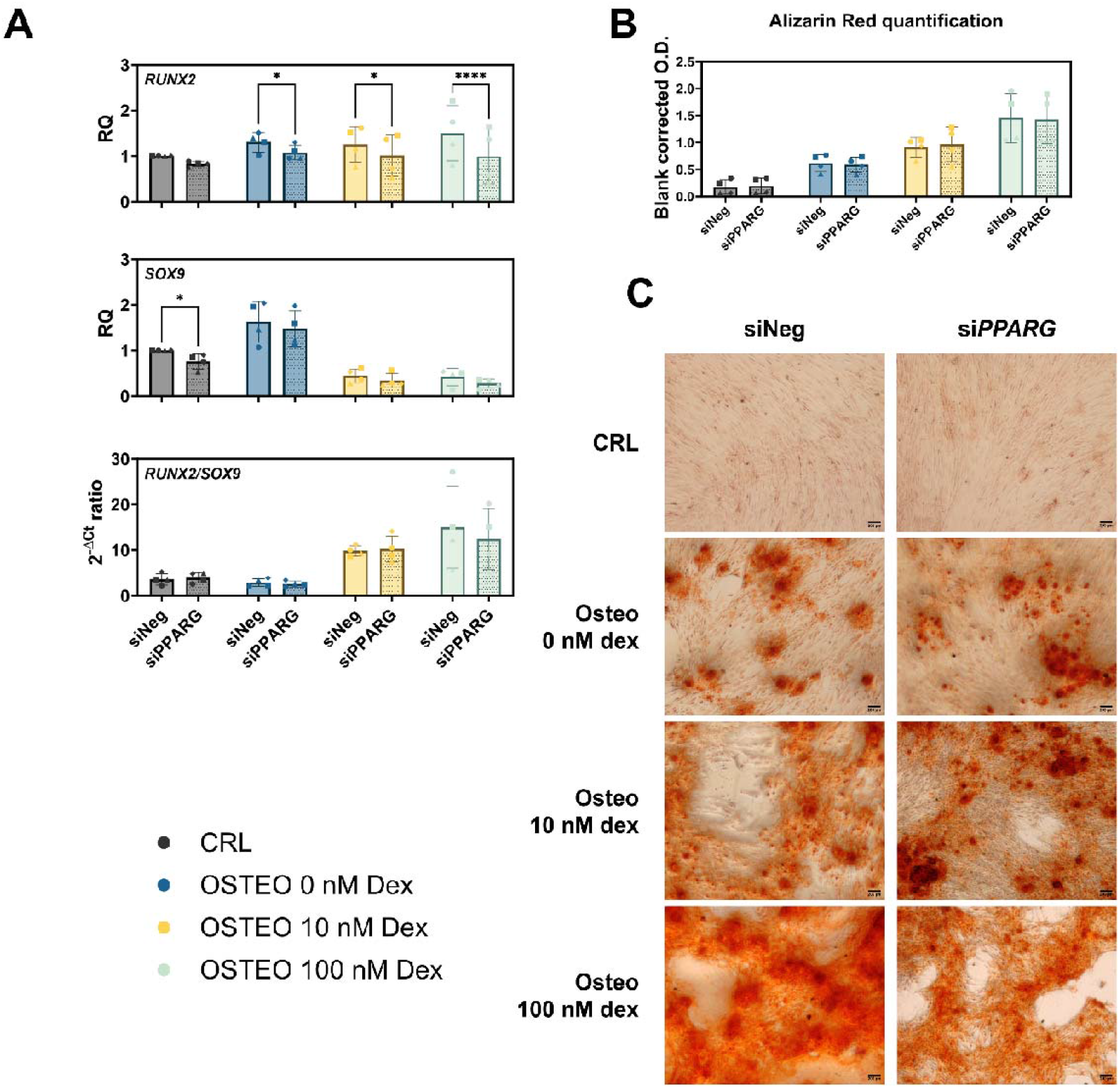
Effect of si*PPARG* treatment on the expression of osteogenic differentiation markers. A) Relative quantification (RQ) of *RUNX2* and *SOX9* expression and *RUNX2*/*SOX9* ratio at day 7. * p < 0.05; **** p<0.0001. B) Alizarin Red quantification from n=4 donors and **C)** representative pictures from 1 donor at day 21 (scale bar = 200 μm). Solid filled bars: siRNA negative control. Dotted bars: treatment with *PPARG*-targeting siRNA.

## DISCUSSION

Dexamethasone is commonly used to induce osteogenic differentiation of human BMSCs *in vitro* (1). However, the effects of dexamethasone on osteogenic differentiation are contradictory, and its overall impact on MSC biology remains not completely understood, prompting reconsideration of its use *in vitro* (6, 23-25). Even though it increases mineral deposition, dexamethasone effects on *in vi*tro osteogenesis could represent artifacts and not recapitulate physiological differentiation mechanisms. The ability of BMSCs to mineralize on a non-osteoconductive surface, such as smooth cell culture plastic in the presence of dexamethasone, supports this hypothesis and may play an important role in the discrepancy between *in vitro* and *in vivo* studies for testing biomaterials for bone regeneration (26).

In an effort to understand the effects of dexamethasone on BMSC osteogenic differentiation to a fuller extent, we previously investigated how different concentrations of the glucocorticoid impact not only osteogenic but also adipogenic and chondrogenic markers. Contrary to common belief, the main transcriptional target of dexamethasone in human BMSCs to induce osteogenic differentiation-like effects is *SOX9*, rather than *RUNX2* (5, 6). SOX9 acts as a dominant-negative regulator of RUNX2 (7), therefore its downregulation allows RUNX2 to exert its effect on its target genes. However, *PPARG* upregulation was an unexpected result, considering the inhibiting role of this transcription factor towards RUNX2 and osteogenic differentiation (16, 27-29).

The results we previously obtained raised several questions and concerns about the use of dexamethasone for inducing osteogenic differentiation *in vitro*. Among the aspects needing elucidation, the mechanism by which dexamethasone leads to *SOX9* downregulation is still unknown. We hypothesized that *PPARG* upregulation by dexamethasone is a main factor regulating *SOX9* expression, meaning that the pro-osteogenic and pro-adipogenic effects of dexamethasone would be closely intertwined. However, PPAR-γ inhibition during dexamethasone-induced osteogenic differentiation did not show the anticipated results. Indeed, *SOX9* expression was not upregulated by PPAR-γ inhibition; rather, we observed no effects on *SOX9* after treatment of human BMSCs with T0070907 or si*PPARG*.

The results from our study are comparable to that by Yu *et al*. (30). In their study, the authors modulated PPAR-γ activity using pharmacological inhibitors (BADGE and GW9662, the latter being closely related to T0070907) or RNA interference by shRNA delivery. Similarly to our results, they found inhibition of adipogenesis, but no improvement of osteogenic differentiation induced by 100 nM dexamethasone. They also did not observe any increase in *RUNX2* expression upon PPAR-γ inhibition. No data about *SOX9* expression is available for comparison. Conversely, other authors reported different results (31-33). Graneli *et al*. conducted an in-depth analysis of the effects of PPARG inhibition using the antagonist GW9662 (32). Their results showed that this inhibition leads to a significant increase in matrix mineralization and a decreased *RUNX2* gene expression. In contrast with our study, in Graneli’s experimental setting the BMSC osteogenic differentiation was induced by the addition of osteogenic medium containing higher concentrations of dexamethasone (1.0 _μ_M), ascorbic acid (45 mM) and _β_ -glycerophosphate (β-GP, 20 mM). Marciano *et al*. observed improved osteogenic differentiation in human BMSCs after PPAR-γ inhibition by pharmacological antagonism (SR2595) or by siRNA treatment (33). Interestingly, dexamethasone was not included in the osteogenic differentiation cocktail used *in vitro*. This suggests that dexamethasone has negative effects on osteogenic differentiation that are independent by PPAR-γ activation, such as osteocalcin inhibition (13, 34, 35). However, it should be noted that we also did not observe an improvement of osteogenic differentiation in our dexamethasone-free controls (CRL and OSTEO 0 nM dex). Another difference between our study and Marciano’s experimental setting is the concentration of β-GP used (10 mM vs 5 mM in our study). β-GP is used as a phosphate source for cell mineralization, as it releases free phosphate ions upon hydrolysis. This is, however, accompanied by the release of glycerol, whose metabolism is regulated by PPAR-γ, at least in adipose tissue (36). Therefore, the effect of different glycerol concentrations might deserve further attention. Other aspects should be carefully considered when comparing the outcomes of PPAR-γ inhibition from different studies. Indeed, there might be species-specific effects (37, 38), a different contribution of the two PPAR-γ isoforms (39, 40), or distinct effects of the inhibitors on PPAR-γ post translational modifications (41).

A logical next step would be animal studies to further validate the functional role of PPAR-γ responses to dexamethasone in bone. However, increasing evidence indicates substantial interspecies differences in drug responses (42, 43). For instance, BMPs consistently induce alkaline phosphatase (ALP) expression and osteogenesis in rodent MSCs, whereas this effect is less predictable in human cells (44, 45). Moreover, BMP ligands and inhibitors often produce contradictory outcomes in human MSCs (46, 47). To address these discrepancies, we propose initial *in vitro* comparative experiments between human and rodent cells to evaluate responses to dexamethasone. Such studies would clarify interspecies variability and strengthen the translational relevance of both *in vitro* and *in vivo* models for bone pathophysiology. Overall, the results of the present study trigger several considerations. Since the effects of dexamethasone on *PPARG* and on *SOX9* expression seem to be independent of each other, it should be possible to uncouple the pro-adipogenic and pro-osteogenic effects of dexamethasone, something that was surprisingly not feasible for BMP-2 mediated osteogenesis (48). Our group has previously investigated the use of a *SOX9*-targeting siRNA (5) in combination with dexamethasone-containing medium. It should be investigated whether *SOX9* knockdown, or other means of SOX9 inhibition, can phenocopy dexamethasone effects on osteogenic differentiation while excluding the adipogenic side effects. Additionally, our results support the hypothesis that dexamethasone effects vary depending on different BMSCs subpopulation (6), something that should be evaluated in future studies with single cell experiments.

## Conclusions

The results presented here suggest that PPAR-γ does not play a role in the repression of *SOX9* and *RUNX2* expression during dexamethasone-induced osteogenic differentiation. On the contrary, targeting PPARG with a silencing RNA reduces RUNX2 expression. On one hand, this finding allows us to explore strategies to separate the pro-osteogenic and pro-adipogenic effects of dexamethasone, potentially improving current protocols for *in vitro* differentiation. However, it also raises further questions about what the true role of dexamethasone, the reliability of our current osteogenic differentiation models, and whether what we observe *in vitro* is an artifact with no parallel to physiological processes.

## List of Abbreviations

α-MEM: Minimum Essential Medium Eagle-Alpha Modification
ADIPOQ: Adiponectin, C1Q And Collagen Domain Containing
bFGF: basic Fibroblast Growth Factor
BMSC: Bone Marrow Derived Mesenchymal Stromal Cells
cDNA: complementary DNA
Dex: dexamethasone-cyclodextrin complex
DMSO: dimethyl sulfoxide
FBS: Foetal Bovine Serum
GR: Glucocorticoid Receptor
LPAR1: Lysophosphatidic Acid Receptor 1
LPL: Lipoprotein Lipase
PPARG: Peroxisome Proliferator Activated Receptor Gamma
rH: relative humidity
RQ: relative quantification
RPLP0: Ribosomal Protein Lateral Stalk Subunit P0
RT-qPCR: Reverse Transcription-quantitative Polymerase Chain Reaction
RUNX2: RUNX Family Transcription Factor 2
siNeg: siRNA negative control
siRNA: small interfering ribonucleic acid
SOX9: SRY-Box Transcription Factor 9

## Declarations

### Ethical approval and consent to participate

The study was conducted in accordance with the Declaration of Helsinki. The Swiss Human Research Act does not apply to research that utilizes anonymized biological material and/or anonymously collected or anonymized health-related data. Therefore, this project did not need to be approved by an ethics committee. Patients’ general consent was obtained, which also covers the anonymization of health-related data and biological material.

Clarification of responsibility: Bern Req-2023-00198.

### Consent for publication

Not applicable.

## Availability of data and materials

The datasets used and/or analysed during the current study are available from the corresponding author on reasonable request.

## Authors’ contributions

Maria Rosa Iaquinta and Carmen Lanzillotti: Data acquisition and analysis, writing – review and editing. Mauro Tognon and Fernanda Martini: Supervision, writing – review and editing. Sonja Häckel: patient selection and management, writing – review and editing.

Martin Stoddart: Conceptualization, supervision, funding acquisition, writing – review and editing.

Elena Della Bella: Conceptualization, methodology, supervision, formal analysis, project administration, funding acquisition, writing – original draft.

All authors read and approved the final manuscript.

## Acknowledgements

The authors thank Zhen Li for providing the T0070907 compound and Cecilia Bärtschi for technical support. The authors declare that they have not used AI-generated work in this manuscript.

## Supplementary file

***Contains* Figure S1**: Representative image of day 21 cultures, treated with 1 _μ_M T00707907, indicating cytotoxicity of the compound at this concentration. **Figure S2:** Representative macroscopic images of Alizarin Red staining from a single donor (n = 1) at day 21. Osteogenic differentiation with different dexamethasone concentration and pharmacological inhibition of PPAR-γ activity with T0070907. Images were acquired using the EVOS2 imaging system (Thermo Fisher) and stitched to provide a complete overview of the well. **Figure S3**: Representative macroscopic images of Alizarin Red staining from a single donor (n = 1) at day 21. Osteogenic differentiation with different dexamethasone concentration and inhibition of PPARG expression with small interfering RNA (siRNA). Images were acquired using the EVOS2 imaging system (Thermo Fisher) and stitched to provide a complete overview of the well.

