## Supplementary file for "Dexamethasone-induced *PPARG* expression in osteogenic differentiation in vitro: impact on *SOX9* and *RUNX2* levels"

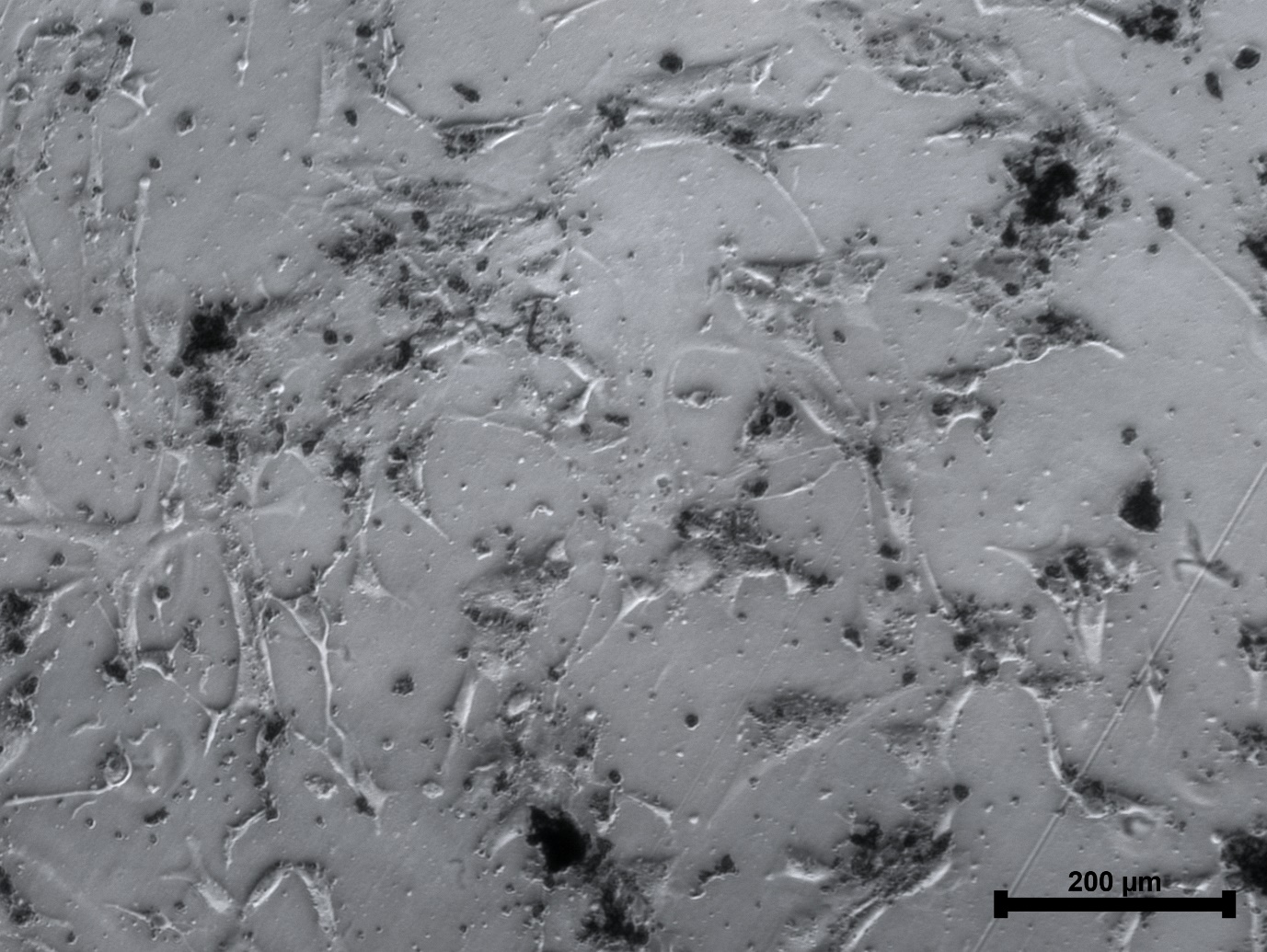

**Figure S1**: Representative image of day 21 cultures, treated with 1 μM T00707907, indicating cytotoxicity of the compound at this concentration.

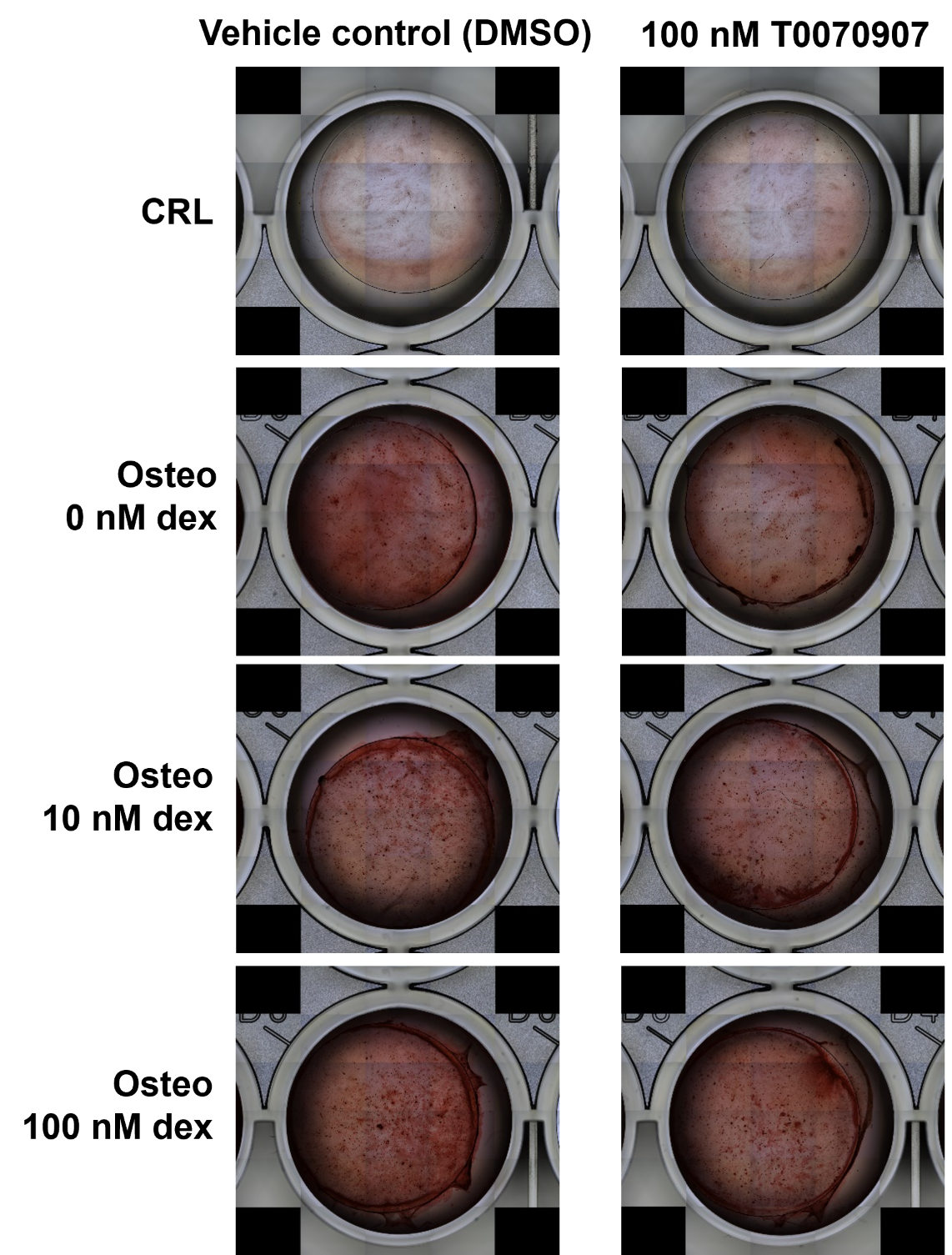

**Figure S2**: Representative macroscopic images of Alizarin Red staining from a single donor (n = 1) at day 21. Osteogenic differentiation with different dexamethasone concentration and pharmacological inhibition of PPAR-γ activity with T0070907. Images were acquired using the EVOS2 imaging system (Thermo Fisher) and stitched to provide a complete overview of the well.

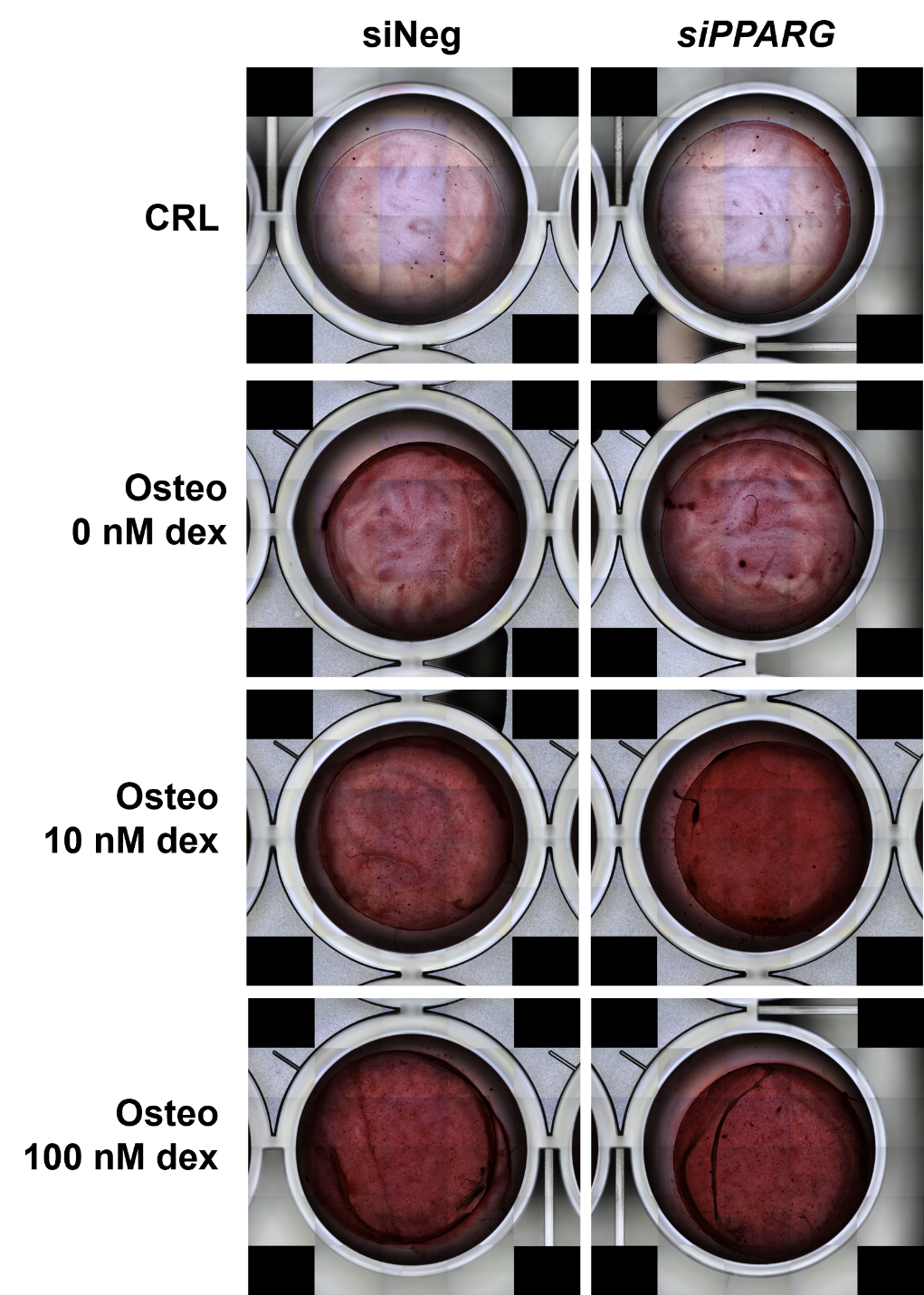

**Figure S3**: Representative macroscopic images of Alizarin Red staining from a single donor (n = 1) at day 21. Osteogenic differentiation with different dexamethasone concentration and inhibition of *PPARG* expression with small interfering RNA (siRNA). Images were acquired using the EVOS2 imaging system (Thermo Fisher) and stitched to provide a complete overview of the well.
